# N-glycome analysis of dried blood spots from different blood preparations and its potential for pre-diabetes and diabetes distinction

**DOI:** 10.64898/2026.08.24.746065

**Authors:** Elham Memarian, Irena Trbojević Akmačić, Ozren Polasek, Gordan Lauc

## Abstract

Dried blood spot (DBS) sampling is becoming a popular alternative to traditional blood sampling approaches, offering advantages such as convenience of collection, transportation, and storage, as well as lower biohazard risk. *N-*glycosylation, a major post-translational modification of proteins associated with numerous biological and pathological functions, is one area of interest for DBS analysis ^1–3^.

In this study, we utilize a protocol for *N-*glycosylation profiling of DBS by ultra-high-performance liquid chromatography based on hydrophilic interactions and fluorescence detection (HILIC-UHPLC-FLR). The protocol includes DBS cutting, protein extraction and enzymatic digestion, labeling with 2-aminobenzamide, followed by cleanup and HILIC-UHPLC-FLR measurement. We compare DBS with plasma and demonstrate the stability of DBS *N-*glycosylation profile when DBS are prepared from fresh blood, frozen whole blood, or a combination of separated frozen blood cells and corresponding frozen plasma. Additionally, we compared DBS *N-*glycans from pre- and diabetic subjects. Fucosylation, bisection, and galactosylation showed a statistically non-significant increasing trend in diabetes, whereas sialylation showed a statistically non-significant decreasing trend in diabetes.

The main advantage of this method is the ability to repurpose samples, which were initially not intended for biomarker *N-*glycan analysis, such as frozen whole blood. Additionally, DBS *N-*glycan profiling is the easier, cheapest and the least invasive approach to conventional plasma in pre-diabetes and diabetes patients’ diagnostics and monitoring.

## Introduction

*N-*glycans are complex sugary molecules that are attached to proteins in the body and play an essential role in a variety of cellular processes ^4^. In recent years, there has been increasing interest in the study of *N-*glycans as biomarkers for a range of diseases, such as diabetes ^5^.

Diabetes is a chronic disease that affects millions of people worldwide, characterized by elevated blood sugar levels that can cause a range of health problems. The most prevalent way of disease diagnosis and monitoring is by measuring glycated hemoglobin (HbA1c), a type of hemoglobin that is formed when glucose attaches to red blood cells ^6,7^. However, HbA1c levels may not always accurately represent a patient’s blood sugar control ^8^, especially in certain populations such as those with anemia or certain genetic disorders ^9,10^. Recent studies have shown that *N-*glycan analysis of plasma samples may provide a more accurate and sensitive biomarker for diabetes compared to HbA1c ^11,12^. In addition, potential associations with type 2 diabetes complications was demonstrated ^13,14^. DBS samples have recently emerged as a promising alternative to traditional blood sampling methods for the screening and monitoring of diabetes ^15,16^. DBS sampling involves collecting a small amount of blood on a filter paper, which is dried and usually stored for later analysis. This method offers several advantages over traditional blood sampling, including easier sample collection, stability during storage and transportation, and decreased risk of biohazard exposure ^15,17,18^.

By analyzing the *N-*glycan profiles of DBS samples, novel biomarkers that can provide insight into disease progression, treatment response, and patient outcomes may be identified. Therefore, this study aimed to undertake the first endeavor by conducting the comprehensive analysis of DBS derived from different blood preparations. This included fresh blood, frozen whole blood, and a combination of separated frozen blood cells and corresponding frozen plasma. The application of this analysis was also performed on samples from individuals with pre-diabetes and type 2 diabetes, marking a significant advancement in this area of research.

## Materials and methods

### Samples

Blood samples were obtained from 40 subjects, divided in three groups. The first group were 32 subjects with pre-diabetes (11 men aged between 37 and 71 years and 21 women aged between 47 and 78 years). The second were 5 subjects with diabetes (2 men aged between 51 and 53, and 3 women aged between 19 and 63). Lastly, the third group consisted of three healthy volunteers, two women aged 32 and 35, and one men aged 52 (Table 1). Samples collected from healthy volunteers were also used for testing different sample collection conditions. The collection of samples was conducted according to the guidelines set out in the ethical approval number 003-08/1 1-03/0005 issued by the university of Split school of medicine ethical board. All participants were from Croatia, and were informed about the study’s objectives before providing their consent. The samples collected were separated into plasma and blood cells, and stored in two separate tubes at a temperature of -20°C.

**Table 1:** Characteristics of the included Croatian samples.

| **Table1: Characteristics of the included Croatian samples.** |  |  |  |
| --- | --- | --- | --- |
|  | **Pre-diabetes** | **Diabetes** | **Healthy** |
| Number of participants, n | 32 | 5 | 3 |
| Sex, n (% male) | 34.4 | 40.0 | 33.3 |
| Age in years, mean | 61.4 | 49.8 | 39.7 |
| Blood glucose concentration, mmol/L, mean (±SD) | 5.4 (±0.5) | 10.1 (±1.9) | - |
| Hemoglobin A1C (HbA1c), %, mean (±SD) | 6.6 (±0.3) | 10.2 (±2.8) | - |

In each plate, two additional positions were filled with phosphate-buffered saline (PBS) solution to serve as blanks_PBS. Two positions were filled with blank_DBS, where the protocol began by cutting a circle from a clean DBS collection card (a card without a blood sample on it). Three positions were allocated for identical plasma samples as technical standards. The plasma sample used as a technical standard was obtained from the Croatian National Institute of Transfusion Medicine after approval by the Ethical Committee of the institute. Additionally, three positions were reserved for pool standard samples (a pool of three plasma samples from healthy individuals). These plasma standards were utilized to assess the accuracy and validity of the measurements. Following the comparison between standard samples and plasma profiles from healthy individuals, the validity of our method was confirmed.

### Condition testing

To study the stability of the samples and to test different conditions, a set of nine DBS from healthy individuals was used. Venous blood was collected from three healthy volunteers, HS1, HS2, and HS3. For each three blood samples obtained from healthy individuals, one ethylenediamine tetraacetic acid (EDTA) blood sample was divided into different parts after incubation at room temperature for 1 hour, except the first condition that involved immediate pipetting onto a collection card without incubation. It is worth mentioning that three replicates of each sample were applied to prepare DBS. After pipetting on collection cards, the blood collection cards were then left to dry overnight. However, all samples from the first healthy sample (HS1) were dried under two different drying times; three hours or overnight at room temperature. After which, all DBSs were placed into plastic bags with desiccants and tightly sealed before being stored at -20 °C until the next step.

For the first condition, freshly collected blood samples were pipetted onto Whatman Schleicher & Schuell bloodstain cards at a volume of 40 μl to prepare DBSs. As the second condition, whole fresh blood samples were separated into plasma and blood cells by centrifuging full blood collected in EDTA tubes, frozen for seven days, then remixed after being defrosted with a ratio of 45% blood cells to 55% plasma (v/v) [whole blood contains around 45% red cells, white cells, and platelets suspended in around 55% blood plasma 19,20]. And DBSs were made from 40 μL of this mixture.

For the third condition, whole collected blood samples were frozen for seven days, defrosted, and then gently mixed in the tube by manually shaking and flipping it upside down. After that, 40 μL of whole blood was pipetted onto the collection card to prepare DBS. The latest condition was only applied to the sample of volunteer HS1. Moreover, separated plasma from these three blood samples was also used for comparison with DBS profile (Table 2) (Figure 1).

**Table 2:** Sample information.

| Table2: Sample information. |  |  |  |  |
| --- | --- | --- | --- | --- |
| Healthy Samples (HS) | Fresh whole blood | Frozen whole blood | Frozen Mix<br>[45%Blood cells+ 55% Plasma (v/v)] | Plasma |
| HS1 | 3 replicates (3hr.)*<br>3 replicates (o. n.)# | 3 replicates (3hr.)<br>3 replicates (o. n.) | 3 replicates (3hr.)<br>3 replicates (o. n.) | 3 replicates |
| HS2 | 3 replicates | - | 3 replicates | 3 replicates |
| HS3 | 3 replicates | - | 3 replicates | - |

\*3hr: Blood spots air-dried for 3 hours at room temperature. #o. n.: Blood spots air-dried over night at room temperature.
\*3hr: Blood spots air-dried for 3 hours at room temperature.

**Figure 1:**
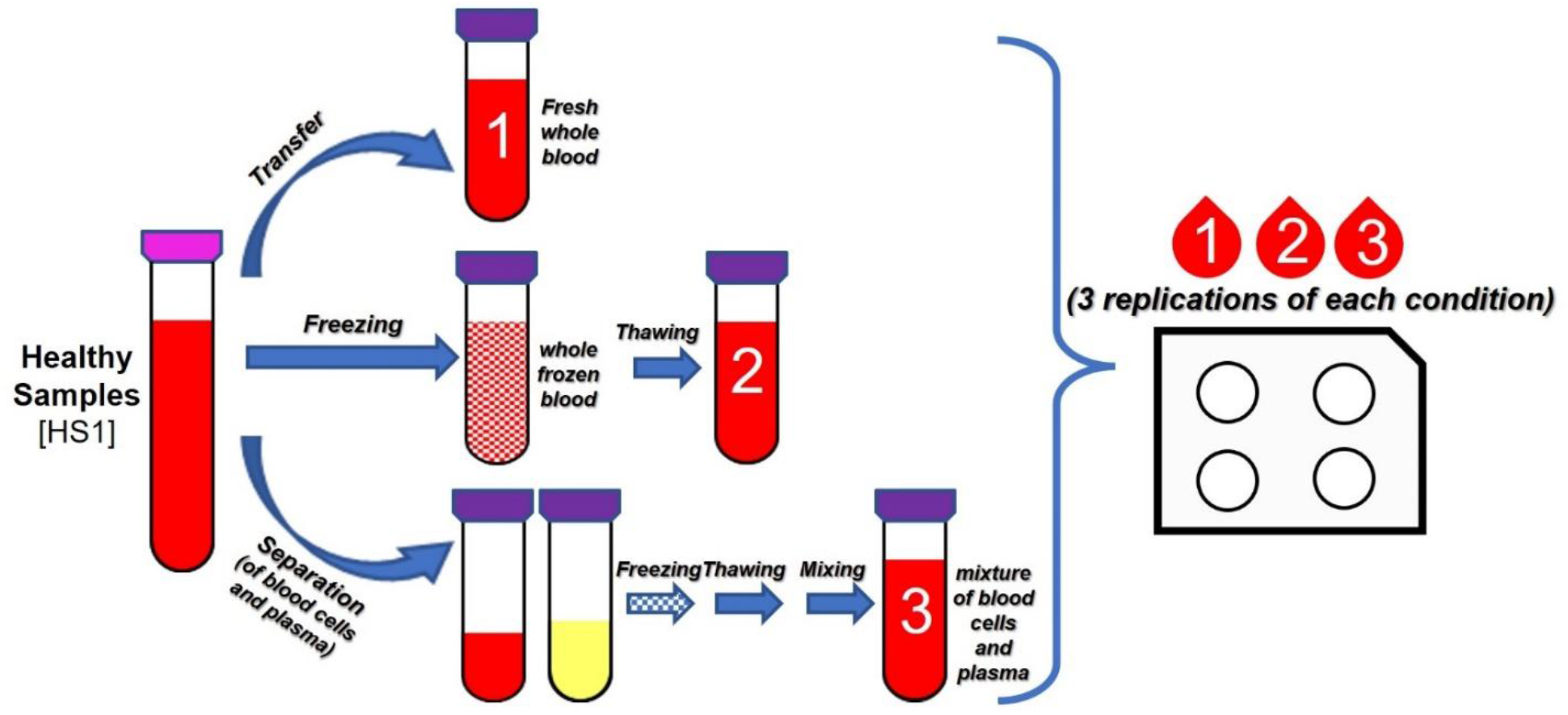
A scheme representing different conditions tested for Healthy Samples (HS).

### N-Glycan release and labeling

A circle with an approximate diameter of 8 mm was cut from each DBS card, and transferred onto a round-bottom plate. For optimal sample preparation throughput, each subsequent step was performed in a 96-well round-bottom plate. Following the addition of 100 μL of ultra-pure water to each well containing a DBS card sample, the next step involved initiating the denaturation process. This was done by the addition of 200 μL of 2% sodium dodecyl sulfate (SDS; w/v; Invitrogen, Carlsbad, CA, USA), and incubating the samples at 65 °C for 10 minutes. Subsequently, 100 μL of 4% Igepal-CA630 (Sigma-Aldrich, St. Louis, MO, USA) and 1.2 U of PNGase F (Promega, Madison, WI, USA) in 100 μL of 5× phosphate-buffered saline (PBS) were added to the samples. The samples were then incubated overnight at 37°C for *N-*glycan release. Following that, samples were dried in a vacuum centrifuge (drying time was around 3 hours). After that, 100 μL ultra-pure water was added into each well and was left on a shaker (room temperature) until the labeling mixture was prepared.

The *N-*glycans were released and labeled with 2-aminobenzamide (2-AB) using a freshly prepared labeling mixture. The mixture was made by dissolving 2-AB (Sigma-Aldrich, St. Louis, MO, USA) in a solution of dimethyl sulfoxide (DMSO; Sigma-Aldrich, St. Louis, MO, USA) and glacial acetic acid (HAc, Merck, Darmstadt, Germany) in a 30% HAc to 70% DMSO ratio. The fresh reducing agent solution, prepared with 2-picoline borane (2-PB, Sigma-Aldrich, St. Louis, MO, USA), was also added to the labeling mixture (for one sample: 0.96 mg of 2-AB, 2.24 mg 2-PB, in 50 μL of 30% HAc in DMSO).

To each *N-*glycan sample in a 96-well plate, 50 μL of the labeling mixture was added, and the plate was sealed with adhesive seal. The mixture was shaken for 10 minutes and then incubated at 65°C for 2 hours. The liquid component of the samples, approximately 150 μL without paper, was then transferred into a new 96-well plate.

To remove free labeling and reducing agents, the samples were cleaned up using a 1 ml AcroPrep wwPTFE 0.2 μm filter plate. The wells were pre-washed three times using 200 μL of 70% ethanol, 200 μL of water, and 200 μL of cold 96% acetonitrile (ACN, Carlo Erba, Cornaredo, Italy), respectively. Next, 1400 μL of cold ACN was added to each sample (2-AB labelled *N-*glycans), mixed by pipetting, and half of this mixture was gently transferred to the wwPTFE plate in two steps using the same tips. The plate was then incubated for two minutes. After two minutes of incubation, the solvent was removed using a vacuum manifold (Millipore Corporation, Billerica, MA, USA). Following this, the second part of the mixture was added, and the plate was further incubated for another two minutes. Subsequently, the second part of the solvent was removed using the same vacuum manifold. The wells were then washed five times using 200 μL of cold ACN/water (96:4, v/v). The glycans were eluted twice, with 90 μL of water for each elution, and the combined eluates were stored at -20°C until usage.

### Hydrophilic interaction liquid chromatography (HILIC)-UHPLC analysis

Fluorescently labeled *N-*glycans underwent separation through hydrophilic interaction liquid chromatography using a Waters Acquity UHPLC instrument. The UHPLC system comprised a quaternary solvent manager, sample manager, and a fluorescence detector. The excitation wavelength was 250 nm, and the emission wavelength was 428 nm. The instrument was controlled by the Empower 3 software, build 3471 (Waters, Milford, MA, USA).

The labeled *N-*glycans were separated using a Waters BEH Glycan chromatography column (150×2.1 mm i.d., 1.7 μm BEH particles). The mobile phase was composed of 100 mM ammonium formate, pH 4.4, as solvent A and ACN as solvent B. During the period of the 25-minute analytical run, a linear gradient ranging from 75% to 53% ACN (v/v) was used at a flow rate of 0.56 ml/min. Before injection, samples were kept at +10 °C, and the separation temperature was +25 °C. The resulting chromatograms were separated into 39 chromatographic peaks, enabling quantification of glycan structures. The percentage of total integrated area for each peak represented the amount of glycans present.

The Empower 3 software was employed to export raw data for 39 glycan structures. From these structures, derived traits that represent the average glycosylation feature shared by different glycan structures were examined. These derived traits were calculated based on the initial glycan traits, as detailed in Supporting Information Table S-1. Standard deviation, coefficient of variation, and other statistical tests (including F-test and t-test) were calculated using Microsoft Excel 2021 (Microsoft 365) and The MathWorks (Natwick, USA).

## Results

### DBS sample preparation for N-glycome profiles

*N-*glycome analysis procedure is based on the previosly published protocol for plasma *N-*glycome analysis^21^. Once the DBS disk is cut and placed into a round-bottom plate, proteins are extracted from the filter paper using a 2% SDS solution, that is identical to the one in the initial plasma protocol. The procedure was adjusted by increasing the total volume of the glycan release reaction and the total volume of the labeling reaction to ensure that the DBS disks are fully immersed in the solution. Furthermore, a drying step and the addition of MQ water to each well of the dried sample prior to labeling step were necessary. Subsequently, the clean-up of the labeled *N-*glycans using AcroPrep wwPTFE, and the measurement using HILIC-UHPLC-FLR was done.

The obtained 2AB-*N-*glycan chromatograms of DBS were comparable to previously reported UHPLC *N-*glycome chromatograms of serum and plasma (chromatograms not shown), in terms of the number of peaks and the general profile ^12,22–24^. It should be noted that all glycan abundances are presented as relative values obtained through normalization based on the summed glycan peak areas. As such, if some signals were to increase, others would decrease.

### Comparison between DBS (venous) with plasma profile

The initial stage of our research focused on assessing the degree of overlap and any potential disparities between the *N-*glycan profiles obtained from DBS and plasma samples. The *N-*glycan profile and peak abundances in DBS chromatogram was highly comparable to the profile generated using our in-house general workflow for plasma sample obtained from a corresponding blood sample (Supporting Information Figure S-1).

The largest difference was around 6% for the most abundant peak. DBS samples exhibited greater signal intensities for glycan profiles at certain peaks, specifically GP1 and GP14, than plasma samples. In contrast, the plasma signal intensities of peaks GP18, GP20, and GP22 were greater than those of DBS. With p-values greater than the predefined cut-off of 0.05, all of the glycan peaks indicated (GP1, GP14, GP18, GP20, and GP22) showed statistically insignificant variations between the DBS samples and plasma.

Based on our results and the outcomes presented in the earlier publication by Vreeker et al. ^15^, which demonstrated that the majority of the released glycans originates from plasma proteins and highlighted that the structural assignment for plasma glycans can be utilized to identify signals within the DBS spectra, the positional and linkage assignments of the *N-*glycans in our study were determined based on UHPLC plasma profile from our previous work ^23^.

### Comparison between DBS obtained from different conditions

Multiple sample preparation and drying periods were investigated for DBS analysis using filter paper cards. Three replicates were measured for each drying period, which were approximately 3 hours and overnight. During both drying periods, samples were completely dried. Our results indicated that glycan profiles obtained from the spots were consistent for both drying conditions. Notably, the DBS samples obtained from the first healthy samples (HS1), which were dried overnight, exhibited comparable profiles to the corresponding plasma samples (Supporting Information Figure S-1).

To investigate different approaches for preparing DBS specifically with regard to the pre- and diabetic samples, we evaluated multiple DBS preparations on filter paper cards. We wanted to explore the possibility of analyzing separated blood cells and corresponding plasma after long-term storage in the form of DBS. . Through our research, we have demonstrated that DBS samples exhibit identical profiles and results regardless of the conditions used to prepare samples. Three replicates were measured from one volunteer (HS1) for each condition, with two different drying periods, for DBS preparation via a mixture of separated frozen-blood cells and corresponding frozen plasma, whole frozen blood, and fresh blood. The glycan profiles obtained from the spots were consistent, as shown in Figure 2. Our results were further validated by measuring DBS preparations via fresh blood and via a mixture of separated frozen-blood cells and corresponding plasma (three replicates of each) from two other volunteers (HS2 and HS3) (Supporting Information Figure S-2 (A, B)).

**Figure 2.**
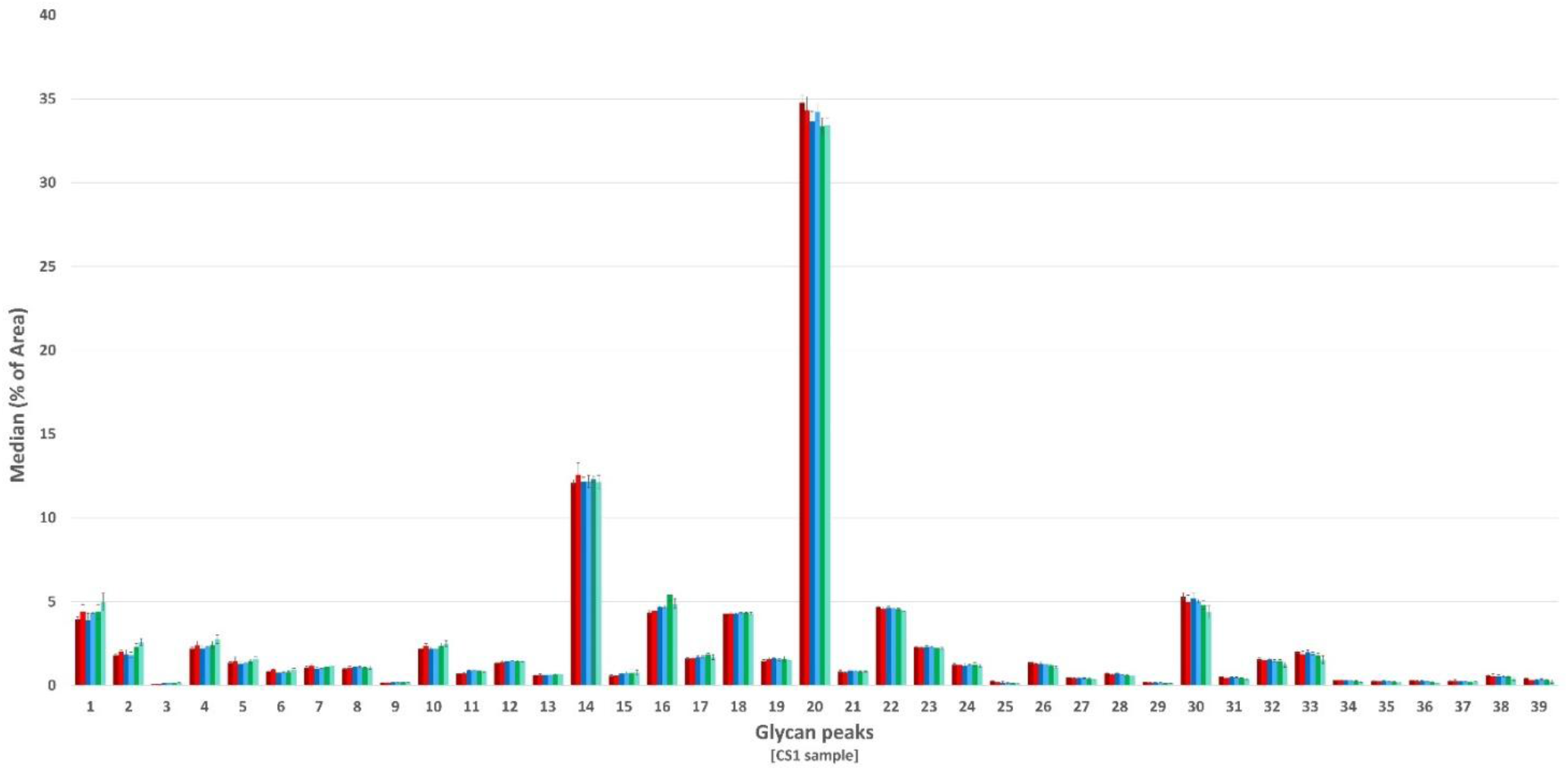
Comparison of N-glycan profiles obtained from DBS samples using three different approaches and two different drying times for a single person (HS1) using HILIC-UHPLC-FLR. The figure displays all glycan peaks, with standard deviation shown as error bars (n = 3). Dark red: DBS obtained from mixture of separated frozen-blood cells and corresponding frozen plasma, drying over night; light red: DBS obtained from a mixture of separated frozen-blood cells and corresponding frozen plasma, drying over 3 hr.; Dark blue: DBS obtained from whole frozen blood, drying over night; Light blue: DBS obtained from whole frozen blood, drying over 3hr.; Dark green: DBS obtained from fresh blood, drying over night; Light green: DBS obtained from fresh blood, drying over 3hr.

### Comparison between cohort DBSs (pre-diabetes versus diabetes)

Mixture of blood cells and its corresponding plasma were used to prepare DBS samples from pre-diabetic and diabetic individuals.

Among 39 directly measured initial glycan traits, after performing t-test and based on the p-values, four studied traits exhibited statistically significant differences, including decreased levels of GP18, and GP29, and increased levels of GP12, and GP22 in diabetic compared to pre-diabetic samples. These associations are displayed in the scatter plot representing the t-test results. The standardized difference between two means was indicated by Hedges’ G. And based on its findings, these significant and non-significant traits exhibited both positive and negative effect sizes, indicating a variety of associational patterns (Figure 3 and Supporting Information Table S-2). To examine differences in directly measured initial glycans, we also searched for significant differences in derived *N-*glycan traits.

**Figure 3.**
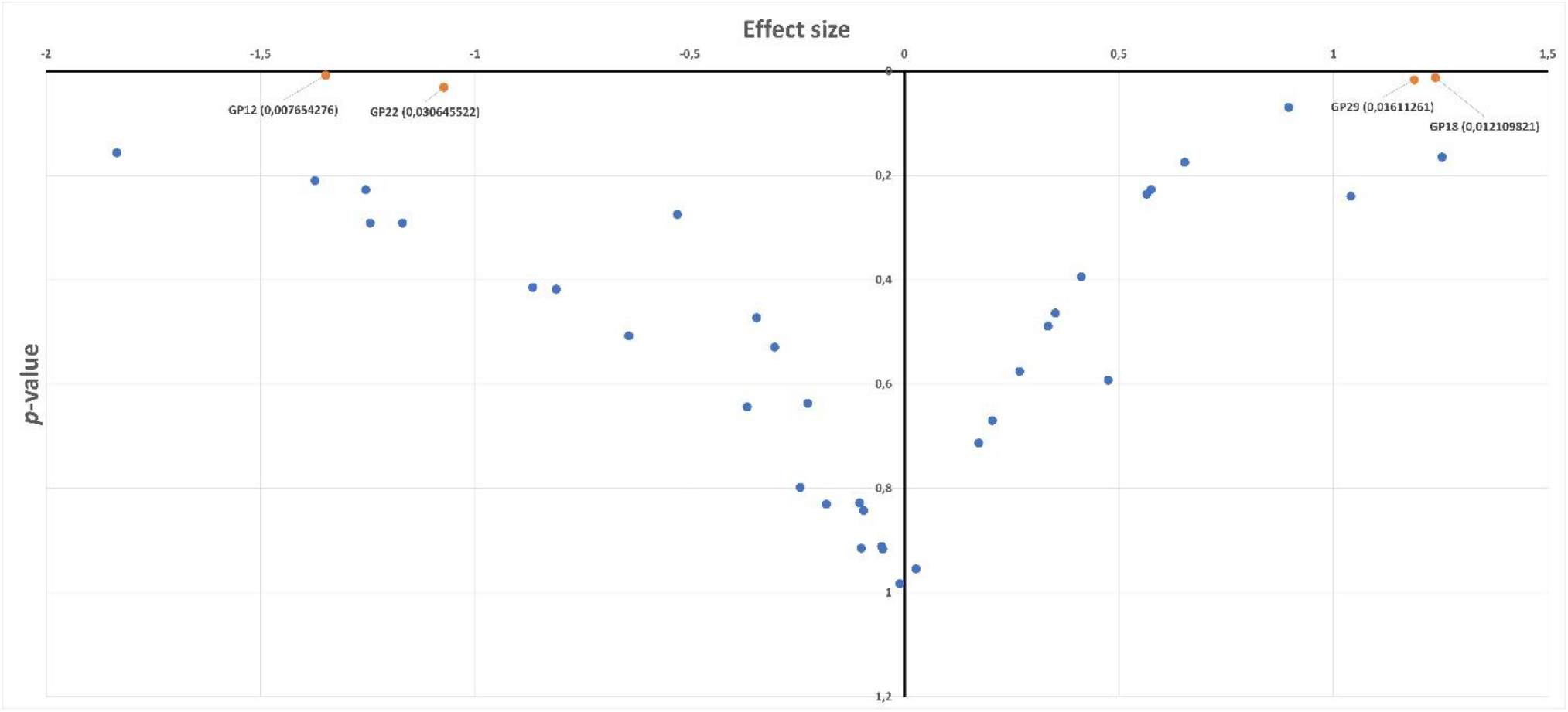
Directly measured N-glycan traits’ p-values plotted against effect size to demonstrate significant associations and patterns of association for pre-diabetes and diabetes samples. The x-axis represents the effect size, calculated as a modified standardized mean difference (Hedges’ g, which is a modified version of Cohen’s d), indicating the magnitude of difference between groups. Hedges’ g provides a more accurate estimate for small sample sizes. The y-axis represents the p-values, indicating the statistical significance of the observed effect sizes. Orange-filled square: Significant association direct traits. Blue-filled square: Non-significant association direct traits.

In diabetic samples, we observed an increase in fucosylation, bisection, and galactosylation, whereas sialylation exhibited the opposite trend (Figure 4). The t-test outcome revealed, however, that these associations were not statistically significant (p-values > 0.05) (Supporting Information Table S-3). This lack of statistical significance can be attributed to the disparity in sample sizes between the pre-diabetic and diabetic groups in our study.

**Figure 4.**
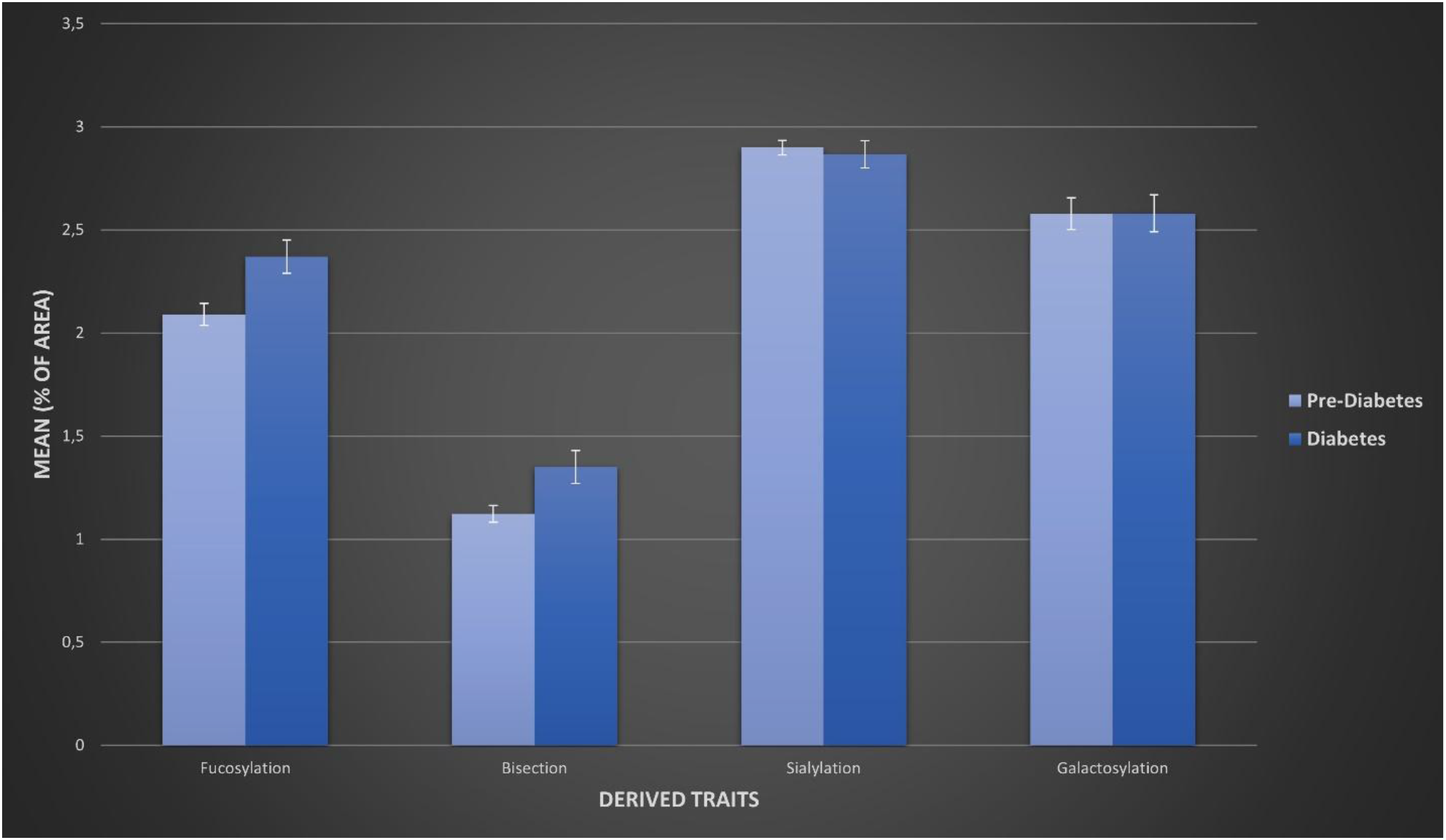
Comparison of derived N-glycan traits obtained from DBS samples for pre-diabetic and diabetic individuals. The figure displays four derived traits. Dark blue: derived traits for DBSs obtained from diabetic samples, light blue: derived traits for DBSs obtained from pre-diabetic samples.

## Discussion

In this study, we present a robust workflow for *N-*glycan profiling using DBSs, which utilizes HILIC-UHPLC-FLR measurement and is well-suited for clinical cohort analysis. Importantly, since glycans are quantified relatively, the analysis is well-suited for the inherently varying volume of blood drops on the filter card. Up to this point, existing research has only examined the analysis of DBS obtained from fresh blood ^15,21^, despite the fact that blood samples are sometimes stored in biobanks as separated frozen-blood cells and corresponding frozen plasma or as frozen whole blood, which may need to be analyzed in DBS format at a later stage. With this in mind, our study aims to address this issue by analyzing DBS samples obtained from differentially stored blood samples and demonstrating its application in distinguishing pre-diabetes and diabetes samples.

Our approach builds upon developed glycomics analysis protocol ^15,21,24^, with several adaptations required to acquire *N-*glycan profiles from a DBS compared to conventional plasma samples, and the most significant modification pertained to the DBS preparation process. Although Gudelj et al. ^21^ previously published *N-*glycome analysis of DBS for age estimation, our method provides a novel approach to the field. The uniqueness of our approach lies in the analysis of DBS samples after separating and individually storing blood cells and plasma. By combining these separate blood fractions, we are able to obtain a DBS sample that is suitable for *N-*glycan analysis, thereby it opens up new possibilities for the research.

UHPLC-FLR analysis was conducted to measure *N-*glycans. The initial step of the experiment involved the investigation of both DBS and plasma samples from a single individual. In addition, using DBS samples obtained from separated frozen-blood cells and corresponding frozen plasma, we were able to analyze both pre-diabetic and diabetic samples, allowing us to distinguish the *N-* glycan profiles of individuals with diabetes from those with pre-diabetes.

The observed differences between the DBS and corresponding plasma samples were limited. Our findings, particularly concerning certain *N-*glycan structures (such as GP16), align with those of previous studies, which demonstrated a higher abundance of IgG-specific glycopeptides in the DBS compared to plasma ^25^. Furthermore, similar to the study conducted by Gerda et al. ^15^, GP14 was found to be slightly more abundant in the DBS compared to plasma.

The DBS profile in our study is consistent with the profiles established in earlier studies regarding DBS and plasma ^21,24^. Our findings, along with previous researches, suggest that the majority of released *N-*glycans in DBS could originate from plasma proteins. In addition, the similarities in glycan profiles between DBS and plasma, as well as the similarity to previous DBS studies performed by our research team, enabled us to propose the glycan structures of each peak in the chromatogram. However, further confirmation by mass spectrometry is required for accurately determining glycan structures in complex chromatograms, such as DBS.

Glycans are recognized for their exceptional chemical stability over time ^17,26,27^. Earlier research has shown the stability of DBS over a long period of time by showing that the relative intensity of 32 N-glycopeptides derived from 14 major N-glycoproteins remained constant throughout storage of DBS samples for 1, 30, 90, and 180 days. ^25^. Additionally, Vreeker et al. reported that glycosylation profiles remained unchanged after storing samples at different temperatures (-80°C, room temperature, and 37°C) for a duration of six weeks ^15^.

The present study investigated the stability of released *N-*glycans using various DBS preparation methods. Our results demonstrated that the glycosylation profiles and relative abundances remained unaffected by the different DBS preparation approaches tested, including fresh blood, whole frozen blood, and separated frozen-blood cells along with their corresponding frozen plasma. This finding illustrates that DBS *N-*glycome analysis can be employed even in situations where collecting DBS from fresh blood samples is difficult or when the original blood samples were not initially collected with the intention of performing DBS experiments.

The study evaluated the short- and long-drying periods of DBS before glycan analysis, and the findings suggested that DBS cards have great potential for facile storage of DBS.

Additionally, our work demonstrates the potential of DBS samples to differentiate pre-diabetic and diabetic *N-*glycan profiles. This is the first report to our knowledge that compares *N-*glycan profiles in pre-diabetes and diabetes. Several studies, however, have studied *N-*glycans in type 2 diabetic samples and compared them to healthy samples. Keser et al. reported an increase in galactosylation, and sialylation, and overall complexity of glycan in diabetic samples, which is associated with an increased risk of type 2 diabetes ^12^. In another study, type 2 diabetes was shown to be associated with a decrease in fucosylation and bisection of diantennary glycans ^28^, and, in the case of IgG, Rudman et al. demonstrated that type 2 diabetes is associated with a decrease in galactosylation and sialylation ^5^.

In our present research, fucosylation, bisection, and galactosylation showed an increasing trend in diabetic samples compared to pre-diabetic samples. In contrast, we observed a decreasing trend of sialylation in diabetic samples compared to pre-diabetic samples. Despite the fact that these results are not statistically significant, the pattern of the results is clear.

Some of our findings, such as the increasing trend of galactosylation in diabetic samples compared to pre-diabetic samples, are consistent with previous research indicating an increased risk of type 2 diabetes associated with increased plasma galactosylation ^12^. Beta-1,4-galactosyltransferases are the enzymes that are responsible for plasma galactosylation, and increased activity of these enzymes in plasma has been associated to both diabetes and aging ^29,30^. However, our findings, particularly regarding sialylation, demonstrate the necessity of using a different point of view when interpreting detailed glycome data from pre-diabetic and diabetic samples. The small sample size employed in the current research may account for these contradictory results. Nevertheless, our study sheds light on the potential of using DBS samples to differentiate between pre-diabetes and diabetes based on their *N-*glycan profiles. And this could lead to the creation of more precise biomarkers and the identification of patients with a greater risk of acquiring diabetes. It is also more convenient and cost-effective approach than traditional glucose blood tests for monitoring pre-diabetic and diabetic patients. Additionally, this approach holds potential as a self-sampling protocol for pre-diabetic and diabetic patients, especially for those who face challenges traveling to a medical facility or require regular monitoring over a longer period ^31,32^. However, given the unequal number of pre-diabetic and diabetic samples used in this study and the relatively small sample size overall, further work on a larger number of samples is warranted to confirm our findings.

It is interesting to note that the DBS from pre-diabetic and diabetic samples for this project were obtained from a mixture of separated frozen-blood cells and corresponding frozen plasma samples, which had been frozen for a couple of years. This opens doors for *N-*glycome analysis in cases when samples have been stored in biobanks for years and shipping the whole blood or separated frozen-blood cells and corresponding frozen plasma is not logistically feasible.

In conclusion, this study presents an analysis of the total *N-*glycome from DBS, focusing on the cases when only frozen blood or separated frozen-blood cells and corresponding frozen plasma are available. Our results show that DBS obtained from fresh blood, frozen whole blood, and a mixture of separated frozen-blood cells and corresponding frozen plasma exhibit similar profiles, thus enabling the *N-*glycan analysis of DBS obtained from various blood sample types.

## Supplementary data

Additional supporting information can be found in the Supporting Information section at the end of the article.

## Author contributions

E.M. contributed to the conception of research question, improved method for glycomics study and performed glycomic analysis of the cohort, processed glycomics raw data, performed data visualization, wrote and reviewed/edited the manuscript. I.T.-A. contributed to the conception of research question and reviewed/edited the manuscript. O.P. contributed to the collection and reviewed/edited the manuscript. G.L. contributed to the conception of research question, reviewed/edited the manuscript, and contributed to the discussion.

### Data availability

The datasets generated during and/or analyzed during the current study are available from the corresponding author upon reasonable request.

### Statements of assistance

The authors thank the study participants and the research staff of the Study in Croatia.

### Guarantor

G.L. is the guarantor of this work and, as such, had full access to all the data in the study and takes responsibility for the integrity of the data and the accuracy of the data analysis.

### Funding

E.M. was supported by funding from the European Union’s Horizon 2020 research and innovation program under the Marie Sklodowska–Curie grant for the project GlySign (contract no. 722095). The study funder was not involved in the design of the study; the collection, analysis, and interpretation of data; writing the report; and did not impose any restrictions regarding the publication of the report.

### Conflict of interest

G.L. is the founder and owner of Genos Ltd, a private research organization that specializes in high-throughput glycomic analysis and has several patents in this field. I.T-A. is an employee of Genos Ltd. E.M. was employed by Genos Ltd at the time of this work; No other potential conflicts of interest relevant to this article were reported.

## References

1. Varki, A. Biological roles of glycans. Glycobiology 27, 3–49 (2017).

2. Esmail, S. & Manolson, M. F. Advances in understanding N-glycosylation structure, function, and regulation in health and disease. Eur J Cell Biol 100, 151186 (2021).

3. Hirata, T. & Kizuka, Y. The Role of Glycosylation in Health and Disease (Chapter:N-Glycosylation). in Advances in Experimental Medicine and Biology 1325 3–24 (Springer, 2021). 10.1007/978-3-030-70115-4_1.

4. Varki A C. R. E. J. et al., editors. N-Glycans (Chapter 9). in Essentials of Glycobiology. (3rd edition); Cold Spring Harbor (NY): Cold Spring Harbor Laboratory Press: New York. (ed. Press, C. S. H. (NY): C. S. H. L.) (2017). doi:10.1101/glycobiology.3e.009.

5. Rudman, N., Gornik, O. & Lauc, G. Altered N-glycosylation profiles as potential biomarkers and drug targets in diabetes. FEBS Lett 593, 1598–1615 (2019).

6. Polonsky, K. S. The Past 200 Years in Diabetes. New England Journal of Medicine 367, 1332–1340 (2012).

7. Odularu, A. T. & Ajibade, P. A. Challenge of diabetes mellitus and researchers’ contributions to its control. Open Chem 19, 614–634 (2021).

8. Youssef D1, E. A. A. J. R. P. A. Fructosamine--an underutilized tool in diabetes management: case report and literature review. Journal of the Tennessee Medical Association 101, 31–33 (2008).

9. Zheng, C. M., Ma, W. Y., Wu, C. C. & Lu, K. C. Glycated albumin in diabetic patients with chronic kidney disease. Clinica Chimica Acta 413, 1555–1561 (2012).

10. Jeffcoate, S. L. Diabetes control and complications: The role of glycated haemoglobin, 25 years on. Diabetic Medicine 21, 657–665 (2004).

11. Tamara, Š. & Lauc, G. N-Glycans and Diabetes. in Reference Module in Life Sciences 1–10 (2021). doi:10.1016/B978-0-12-821618-7.00001-8.

12. Keser, T. et al. Increased plasma N-glycome complexity is associated with higher risk of type 2 diabetes. Diabetologia 60, 2352–2360 (2017).

13. Memarian, E. et al. Plasma protein N-glycosylation is associated with cardiovascular disease, nephropathy, and retinopathy in type 2 diabetes . BMJ Open Diabetes Res Care 9, e002345 (2021).

14. Memarian, E. et al. IgG N-glycans are associated with prevalent and incident complications of type 2 diabetes. Diabetes Metab Res Rev 39, e3685 (2023).

15. Vreeker, G. C. M. et al. Dried blood spot N-glycome analysis by MALDI mass spectrometry. Talanta 205, 120104 (2019).

16. Plamen A. Demirev. Dried Blood Spots: Analysis and Applications. Anal. Chem. 85, 779–789 (2013).

17. Ruhaak, L. R.; Miyamoto, S.; Kelly, K.; Lebrilla, C. B. N-Glycan Profiling of Dried Blood Spots. Anal Chem 84, 396–402 (2012).

18. Skogvold, H. B., Rootwelt, H., Reubsaet, L., Elgstøen, K. B. P. & Wilson, S. R. Dried blood spot analysis with liquid chromatography and mass spectrometry: Trends in clinical chemistry. Journal of Separation Science 46, 202300210 (2023).

19. visiblebody-Learn Site. Overview of Blood [Date accessed: 2023-08-10]. visiblebody https://www.visiblebody.com/learn/biology/blood-cells/blood-overview.

20. Mathew J, S. P. V. M. Physiology, Blood Plasma. [Updated 2023 Apr 24]. (In: StatPearls [Internet]. Treasure Island (FL): StatPearls Publishing; (Available from: https://www.ncbi.nlm.nih.gov/books/NBK531504/), 2023.

21. Gudelj, I. et al. Estimation of human age using N-glycan profiles from bloodstains. Int J Legal Med 129, 955–961 (2015).

22. Zaytseva, O. O. et al. Heritability of Human Plasma N-Glycome. J Proteome Res 19, 85–91 (2020).

23. Adua, E. et al. High throughput profiling of whole plasma N-glycans in type II diabetes mellitus patients and healthy individuals: A perspective from a Ghanaian population. Arch Biochem Biophys 661, 10–21 (2019).

24. Trbojević Akmačić, I. et al. High-throughput glycomics: Optimization of sample preparation. Biochemistry (Moscow) 80, 934–942 (2015).

25. Choi, N. Y. et al. Direct analysis of site-specific N-glycopeptides of serological proteins in dried blood spot samples. Anal Bioanal Chem 409, 4971–4981 (2017).

26. Gornik, O. et al. Stability of N-glycan profiles in human plasma. Glycobiology (2009) doi:10.1093/glycob/cwp134.

27. Simunovic, J. et al. Comprehensive N-glycosylation analysis of immunoglobulin G from dried blood spots. Glycobiology 29, 817–821 (2019).

28. Dotz, V. et al. Plasma protein N-glycan signatures of type 2 diabetes. Biochim Biophys Acta Gen Subj 1862, 2613–2622 (2018).

29. Lee, L. P. K. et al. Serum UDP-Galactose.” Glycoprotein Galactosyltransferase in Diabetics with Microangiopathy. Clin. Biochem 10 111–117 (1977).

30. Catera, M. et al. Identification of novel plasma glycosylation-associated markers of aging. Oncotarget 7, 7455 (2016).

31. Taber, J. M., Leyva, B. & Persoskie, A. Why do People Avoid Medical Care? A Qualitative Study Using National Data. J Gen Intern Med 30, 290–297 (2015).

32. Anderson, L. Six decades searching for meaning in the proteome. J Proteomics 107, 24–30 (2014).

